# Monitoring seagrass meadow health using coastal bird fecal fatty acid biomarkers

**DOI:** 10.64898/2026.09.01.748668

**Authors:** Christopher Mulligan, David A. Gold

## Abstract

Lipids have the potential to trace trophic interactions through an ecosystem, but their use has been largely limited to predator/prey-scale studies. Carnivore fecal lipids offer an unexplored way to strategically monitor the health and spatial distribution of ecosystems that are difficult to access. This hypothesis is explored here in seagrass meadows, which are a globally significant long-term carbon storage reservoir. This study tested the hypothesis that lipids can be used to passively monitor seagrass meadow health by tracing fatty acids transferred up the food chain and onto nearby beaches via coastal bird stool. By comparing the lipids produced by seagrasses and those made by diverse microbial eukaryotes, such as diatoms and dinoflagellates, our analysis could distinguish between healthy and collapsed meadows in a pilot dataset. A metabolite that is produced by the seagrass wasting disease pathogen was also recovered in our data, providing a potential diagnostic for seagrass health. The advantages to this non-invasive monitoring approach using lipids are that beaches are far more accessible than seagrass meadows, and birds spatially ‘sample’ a larger area than researchers could do efficiently. Although this pilot study was limited in samples and field sites, our results provide compelling reasons to pursue carnivorous fecal dietary lipids as a source of ecosystem data.

## Introduction

Seagrass ecosystems are incredibly important on a global scale but are vulnerable to disturbance. Seagrass meadows provide hunting grounds for many animals including seals and birds, act as nursery habitats for several species of fish and invertebrates, stabilize coastal erosion, and play a significant role in mitigating climate change both from their efficient carbon sequestration and by buffering ocean acidification (Hyndes et al. 2018; Unsworth et al. 2019; Ricart et al. 2021). Despite the multitude of benefits these ecosystems provide, there are many threats to meadows globally (Unsworth et al. 2019; 2022). Like many ecosystems, seagrass meadows are facing a combination of risks, including increasing seawater temperatures, pollution and runoff, and wasting disease infections caused by the pathogen *Labyrinthula*. Collectively these threats can result in the collapse of seagrass meadows, leaving either bare mudflats or algal/biofilm dominated communities in their wake (Unsworth et al. 2019; 2022). The loss of these important ecosystems leads to a loss of carbon storage and all of the other benefits they provide. There are many long-term seagrass observation stations across the globe, most of which fall under the SeagrassWatch and SeagrassNET organizations. Monitoring can include many variables, including area coverage, shoot density, visual ID of lesions from wasting disease, C/N/P elemental ratios in the shoots, and environmental information such as light, depth, and temperature (Unsworth et al. 2014). Some typical methods for meadow health and extent include intertidal walking and plant sampling, freediving, towed videos, sonar, and helicopter (Unsworth et al. 2014). While these techniques are the field standard for monitoring seagrass meadows, they can be timely, exhausting, and expensive to carry out for long term records, especially at short intervals. Tracking seagrass carbon sequestration is typically done with isotopes from the total carbon pool, but some of those individual molecules are very biologically specific and that information is often lost with isotopic analyses (Ward et al. 2021). Given the importance of seagrass meadows and the difficulty of reaching them, additional monitoring methods would be scientifically valuable.

A possible alternative approach to monitoring seagrass meadows involves tracing their lipids through the coastal ecosystem. Fatty acids (or FAs) are a common lipid class in cell membranes composed of hydrocarbon chains, usually 14 to 24 carbons long, with a terminal carboxyl headgroup. FA compound notation is structured using the following notation: the total number of carbons in the compound, the number of saturations, and the position of the first saturation starting from the carboxyl end. For example, 22:6n-3 is 22 carbons long with six double bonds beginning at the third carbon from the carboxyl end. FAs are used in cells for membrane stability and permeability, energy storage, cell signaling, and as the building blocks for more complex lipids (de Carvalho and Caramujo 2018). Lipid biomarkers have been used in many studies to track ecological interactions by taking advantage of taxa-specific compounds that get incorporated into the tissues of consumers, thus bypassing the limitations of observation or gut contents (Jordan 2005; Nielsen et al. 2018; Galloway and Budge 2020; Jardine et al. 2020). FAs are particularly useful for distinguishing bacteria, phytoplankton, and algal prey (Jardine et al. 2020, Kelly and Scheibling 2012). Since the FAs of autotrophs are chemically similar to the consumer’s FAs they require relatively little (and usually predictable) modification before incorporation into cell membranes (de Carvalho and Caramujo 2018). While there is a wealth of information that could be learned from seagrass lipidomics, several key FAs might be particularly useful to monitor their health (Sbrizzi et al. 2024). Seagrasses can be characterized using 18:2n-6, 18:3n-3, and 28:0 FAs; while these are typical FAs in flowering plants, they can be useful when paired with additional lipid analyses that calculate terrestrial versus marine biomass (Dalsgaard et al. 2003; Alfaro et al. 2006; Colombo et al. 2017; Dahl et al. 2025). Long chain FAs (LCFAs), particularly those longer than 24 carbons, are also used to identify plant leaf waxes from flowering plants, including seagrasses (McClymont et al. 2023). In contrast, green algae such as sea lettuce (*Ulva* spp.), which often dominate an area after seagrass meadows collapse, can be identified by diverse n-6 polyunsaturated FAs (PUFAs) (Galloway et al. 2012). Many other clades found in coastal ecosystems, such as kelps and other brown algae, diatoms, and golden algae, also have distinct FAs and can be reliably identified using lipids (Galloway et al. 2012). One commonly used class of FAs are omega FAs, also called essential FAs, which cannot be synthesized by most animals but are critical for their musculature and nervous systems. The omega fatty acid docosahexaenoic acid, or DHA (22:6n-3) is highly associated with marine ecosystems but is not produced by macroalgae or seagrasses, instead being biosynthesized primarily by microbial eukaryotes such as diatoms, dinoflagellates, cryptomonads, and notably the eelgrass pathogen *Labyrinthula zosterae* (Kharlamenko et al. 2001; Dalsgaard et al. 2003; Veloza et al. 2006; Chu et al. 2008; Coelho et al. 2011; Galloway et al. 2012; Peltomaa et al. 2017; Yoshioka et al. 2019). DHA is highly retained trophically, with nearly a 1:1 transfer rate between prey and predators (Jardine et al. 2020). These major groups of autotrophs that form the foundation of marine ecosystems have distinct lipid profiles, making lipidomics a powerful tool to study offshore communities.

It has long been recognized that animals take on a FA signature reflecting their diet, or their prey’s diet, allowing researchers to estimate the trophic linkages using lipid tracers (Nichols et al. 1986; Alfaro et al. 2006; Budge et al. 2006; Coelho et al. 2011; Kelly and Scheibling 2012; Guerrero and Rogers 2020). The combined facts that seagrass meadows have distinct FA profiles and that FAs can be transferred up the food web means that nearshore birds have the potential to retain FA biomarkers from these offshore ecosystems. In this system (illustrated in Figure 1) the lipids of seagrasses and other autotrophs get taken up by primary consumers (such as snails and amphipods) and move up the food chain into generalist predators. Each animal in this series will typically esterify their dietary lipids to remove the headgroup, then either incorporate them into their cell membranes or store them for energy in compounds like triacylglycerides (Giese 1966; Griminger 1986; Williams and Buck 2010). Once the marine animals are eaten by a bird, the prey’s tissues and stomach contents are broken down and the FAs are released once again. While the majority of FAs are reabsorbed through the intestinal lining, carnivorous birds have short digestive tracts and relatively quick food retention times, meaning a portion of these diagnostic lipids are potentially passed out as waste and amenable for research (Williams and Buck 2010; Twining et al. 2021). FAs have been used extensively in birds, using tissues such as feathers, eggs, blood plasma, and adipose samples to reconstruct their diets (Williams and Buck 2010). Marine and coastal birds serve as important sentinel species, both their presence and health can be used as proxies for the larger ecosystem’s condition (Boersma 2008; Grove et al. 2009; Wolf et al. 2010; Quinn et al. 2017; Mathot et al. 2019). One important use of birds is for ecotoxicology monitoring; as predators they biomagnify pollutants such as lead and microplastics that propagate up the food chain (Burger and Gochfeld 2004; Grove et al. 2009; Athira et al. 2024; Montgomery et al. 2026). Several recent studies have developed other novel techniques using data from wildlife to monitor ecosystems that are difficult to access by taking advantage of an animal’s feeding habits and range coverage (Siegenthaler et al. 2019; Shao et al. 2021; Ji et al. 2022; Hartig et al. 2024; Beltran et al. 2025).

**Fig 1:**
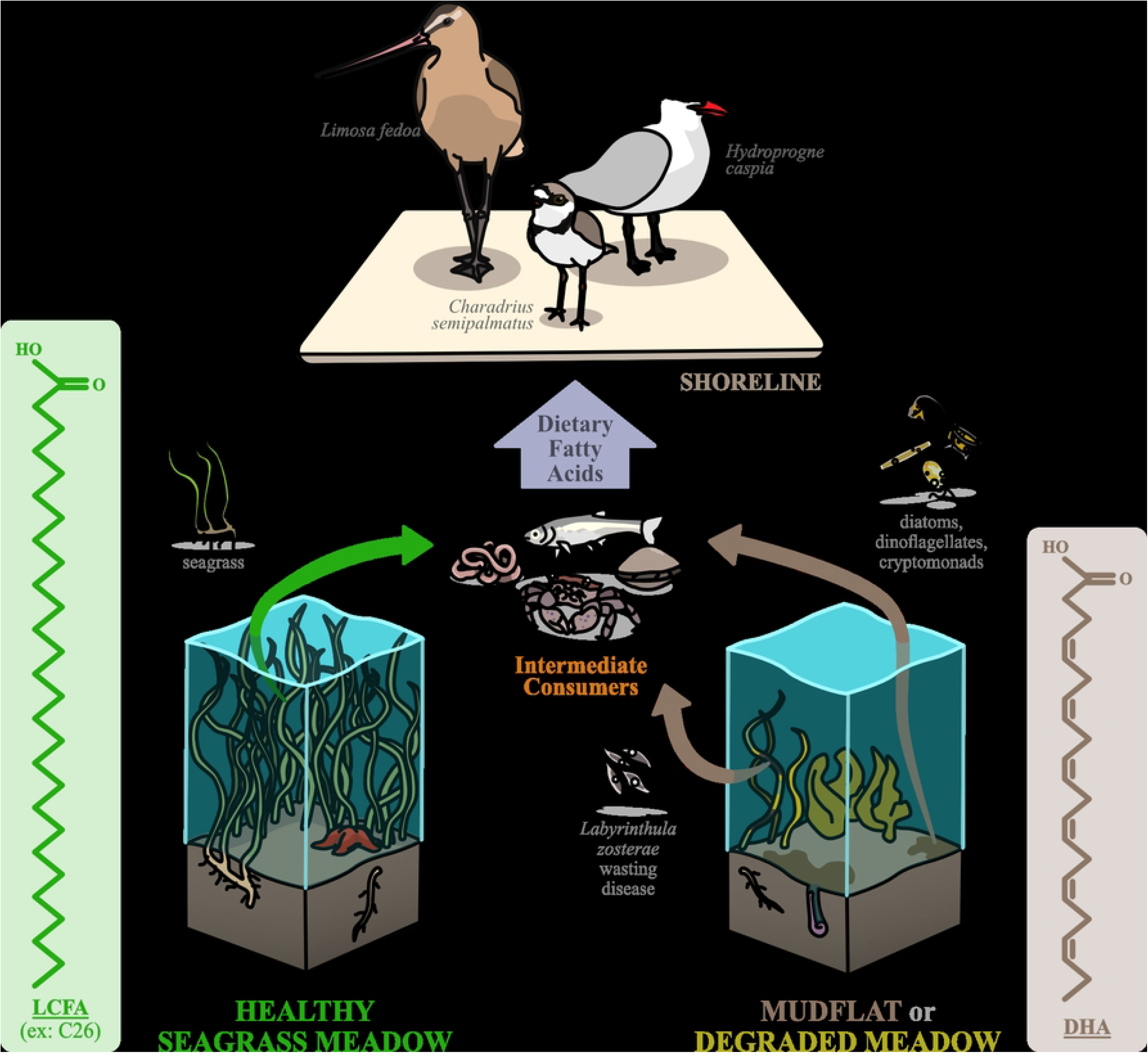
Transfer of fatty acids along trophic relationships proposed in this study, traveling from autotrophs, namely long-chain fatty acids (LCFAs) produced by seagrass shown in green and docosahexaenoic acid (DHA) rich microbial eukaryotes (shown in brown), up through intermediate prey species and ultimately deposited onto the shoreline by birds.

This pilot study explored if coastal bird fecal lipids correlate to the local offshore ecosystems, a mudflat and a seagrass meadow, as confirmed through field observations. Our hypothesis was that FAs that were originally biosynthesized by seagrass and microbial eukaryotes can be traced up the marine food web and ultimately recovered in the bird feces on the beach. By leveraging shorebird ecology with lipid biomonitoring principles, it may be possible to employ wildlife to do a majority of the fieldwork for tracking seagrass meadow health without researchers ever leaving the shoreline.

## Material and methods

### Sample Collection

The birds targeted in this project were carnivorous shorebirds in the order Charadriiformes (plovers and relatives), with additional samples from Anseriformes (ducks and their relatives) and Ardeidae (herons and relatives). Droppings from seagrass grazers and biofilm-specialists such as geese and small sandpipers were largely avoided since these birds prefer consuming autotrophs and therefore would have a biased signal against nonvegetated mudflats (Valentine and Heck 1999; Kuwae et al. 2012). In contrast, medium and large carnivorous shorebirds (e.g. scoters, avocet, plovers) consume a wide range of generalist invertebrates or fish that are present in all nearshore ecosystem types (Angarita-Báez and Carlos 2023; Correia et al. 2023).

Samples were collected from two field sites between May and July 2025. Millerton Point in Tomales Bay, California (on public land approximately 38.314573, -123.056826) was chosen as a healthy seagrass site since wasting disease is uncommon until much later in the summer, based on many years of continuous monitoring by several groups at the UC Davis Bodega Marine Laboratory (BML). Gaffney Point in Bodega Harbor, California (located partially on the Bodega Marine Reserve and on public land at approximately 38.105780, -122.849698) is a tidal mudflat with very small declining patches of seagrass that consistently have wasting disease lesions; for simplicity this will be referred to as the “mudflat” ecosystem. No collection permits were required since fecal materials are not covered by state or federal restrictions and the collection was only done after the animals left the area so as not to disturb them or alter their behaviors. Samples were collected during the day under overcast to sunny conditions when the birds were most active. Sample collection was largely timed with mid- to low-level tides, when enough area was exposed for foraging but the tide was high enough to keep the birds close to the main shoreline. A rising tide will push the birds closer to the shore but comes with increased risk of losing samples to the ocean. Additional samples were collected without having observed the producing species but were ultimately omitted from most statistical analyses (i.e. PCA and ANOVA).

All field work was conducted with appropriate aseptic techniques and PPE out of caution for a regional outbreak of H5N1 Highly Pathogenic Avian Influenza (HPAI), of which many of this project’s target species are reservoir hosts (CDC, 2024). Birds were monitored along the shoreline, and fresh fecal samples were collected after observing the deposit, identifying the species, and monitoring for visual symptoms of HPAI. Stools were sampled in a way that minimized urates and excluded any environmental sediment, even at the expense of stool material. When cecal droppings were found they were ignored since it would likely reflect the bird’s long-term diet and not the last meal. Samples were collected with a sterile plastic scoop into pre-weighed and labeled borosilicate 20 mL vials containing ∼10 mL of pre-aliquoted ethanol (EtOH). A square of furnace-baked aluminum foil was placed between the glass and vial cap. These vials were gently mixed to get the stool fully submerged before continuing with field work. Vial exteriors were thoroughly cleaned with Super Sani-Cloth® sanitizing wipes and allowed to air dry for at least 1 minute. All waste was double bagged, with the first bag being washed with sanitizing wipes and air dried before adding the second bag, which was also wiped down. All PPE (goggles, waders, and boots) were rinsed with fresh water both on location and at BML, then treated with wipes after each field day and allowed to air dry. A field control was collected by opening a vial in the intertidal zone of Gaffney Point, closing it, and storing/processing it with the other samples. Control samples of healthy seagrass (*Zostera marina*) and seagrass lesions infected with wasting disease (*Labyrinthula zosterae*) were collected from mesocosms grown at BML and stored in EtOH, courtesy of Serina Moheed.

### Data Analysis

Samples were stored at room temperature in a fume hood, then submitted to West Coast Metabolomics Center for extraction and analysis by ultra high performance liquid chromatography (UHP-LC) following the methods outlined in Matyash et al. (2008). The relative abundances of free FAs (FFAs) were normalized as a sum of the total FFAs to make the data directly comparable across samples. Analysis was limited to FFAs rather than intact lipids (i.e. those with a headgroup or part of larger storage molecules like triacylglycerides) for three reasons: 1) feces is chemically complex and fewer unique molecules meant a simpler analysis, 2) intact lipids typically represent living biomass, and we wished to exclude both the bird gut microbiome or any potential environmental contamination, and 3) dietary intact lipids may lose their biological specificity after being passed through multiple consumers, making the analysis of headgroup-specific affinities contentious. Principal components analysis (PCA, via the R function prcomp in the Stats package v4.5.2) was performed on the full UHP-LC results (Supplementary Figure 1) as well as the FFAs. Analysis of variance (ANOVA, via the R function aov also from the Stats package v4.5.2) tests using 8 different lipid classes and proxies (see **Table 1**) were completed to determine the significance of sample location. To assess the goodness-of-fit of the high p value models, separate ANOVAs were done on the linear regression of those models to report the r^2^ value.

**Table 1:** Results of each ANOVA test. SFAs = saturated free fatty acids (FFAs), MUFAs = monounsaturated FFAs, PUFAs = polyunsaturated FFAs, LCFAs = long chain FFAs (24:0 + 24:1n-9 + 25:0 + 26:0 + 28:0 + 29:0 + 30:0). The FFAs for seagrass biomarkers were 18:2n-6 + 18:3n-3 + 28:0. A ratio of terrestrial (18:2n-6 + 18:3n-3) vs marine (20:4n-6 + 20:5n-3 + 22:6n- 3) PUFAs was calculated following Colombo et al. (2017). An asterisk* denotes statistically significant P values below 0.05.

| Test | Mean Sq | F value | P value | Residuals Mean Sq |
| --- | --- | --- | --- | --- |
| All FFAs | 9.69E-28 | 1.404 | 0.257 | 6.90E-28 |
| SFAs (0 saturations) | 706.9 | 1.84 | 0.198 | 384.1 |
| MUFAs (1 saturation) | 204.2 | 1.006 | 0.334 | 203 |
| PUFAs (>2 saturations) | 1671 | 3.664 | 0.0779 | 456 |
| LCFAs (>C24) | 101.99 | 3.632 | 0.079 | 28.08 |
| Seagrass biomarkers | 0.222 | 0.063 | 0.805 | 3.499 |
| DHA / (DHA + LCFA) | 0.5058 | 7.747 | **0.0155\*** | 0.0653 |
| Terrestrial vs marine PUFAs | 0.004783 | 0.969 | 0.343 | 0.004938 |

## Results

A total of 24 stool samples were collected: nine from Millerton Point and 15 from Gaffney Point. Samples were collected from seven species: great egret (*Ardea alba*, n=2), sandpiper (*Calidris* spp., n=1), semipalmated plover (*Charadrius semipalmatus*, n=2), Caspian tern (*Hydroprogne caspia*, n=4), marbled godwit (*Limosa fedoa*, n=4), Hudsonian whimbrel (*Numenius hudsonicus*, n=1), and a willet (*Tringa semipalmata*, n=1). It is worth noting that, unfortunately, there were no samples collected from the same species at both sites. Nine additional samples were opportunistically collected without observing the individual that produced them. UHP-LC analysis returned 4,067 compounds of which 1,202 were characterized (Supplementary File 1) and 38 were FFAs (seen in **Figure 2**, with each as a proportion of the total FFAs; Supplementary File 2). Nomenclature for FFAs was standardized using both the Metabolomics Workbench (https://workbench.sdsc.edu/databases/refmet) and PubChem (https://pubchem.ncbi.nlm.nih.gov/) databases. All code and data can be found at this GitHub repository: https://github.com/DavidGoldLab/2026_Seagrass_Monitoring/tree/main.

**Fig 2:**
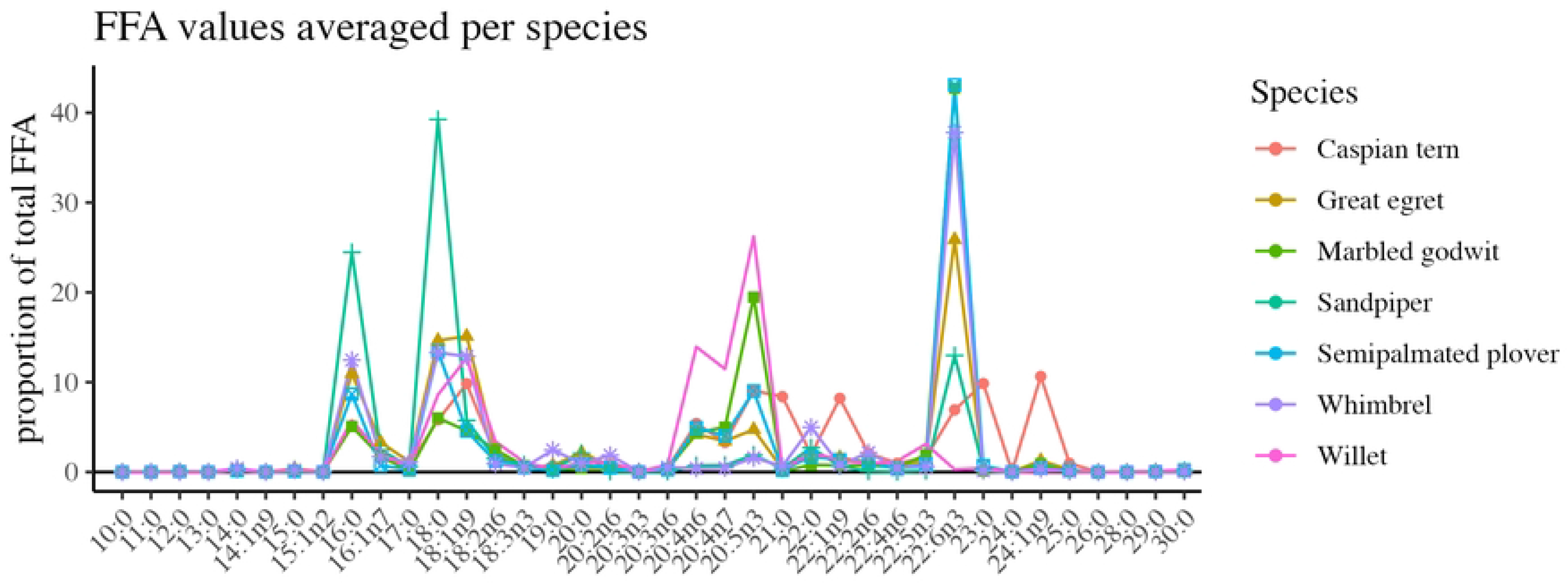
Average fecal free fatty acids (FFA) proportion for each bird species.

PCA results suggests that healthy and collapsed seagrass meadows can be distinguished by their FFA profiles (**Figure 3**). Principal component (PC) 1 and PC2 (Figure 3A-B), captured 55.6% of the variation and separated the two ecosystems, while PC2 and PC3 (Figure 3C-D), captured 34.65% of the variation and separated the bird species. PC2 differentiates the samples largely along axes of 16:0 + 18:0 and 21:0 + 23:0, while PC1 and PC3 were most strongly associated with 22:3n-6 and 20:5n-3, respectively. Following these results, barplots (**Figure 4**) were created showing the normalized ratio of DHA versus LCFAs (24:0 + 24:1n-9 + 25:0 + 26:0 + 28:0 + 29:0 + 30:0) for both fecal samples (including unknown species) and seagrass tissue controls from BML mesocosms. The healthy meadow’s lowest values (Figure 4B) were associated with species observed feeding in the seagrass (i.e. the willet was foraging in an intertidal seagrass meadow and the terns were seen feeding for most of the day in a deep channel of subtidal grass) while higher values came from species feeding closer to shore in marginal nonvegetated mud areas (i.e. the plovers and sandpiper). The sandpiper values may also reflect the influence of biofilms in their diet, potentially explaining why their PCA values are so far removed from the other species (Kuwae et al. 2012; Schnurr et al. 2020).

**Figure 3:**
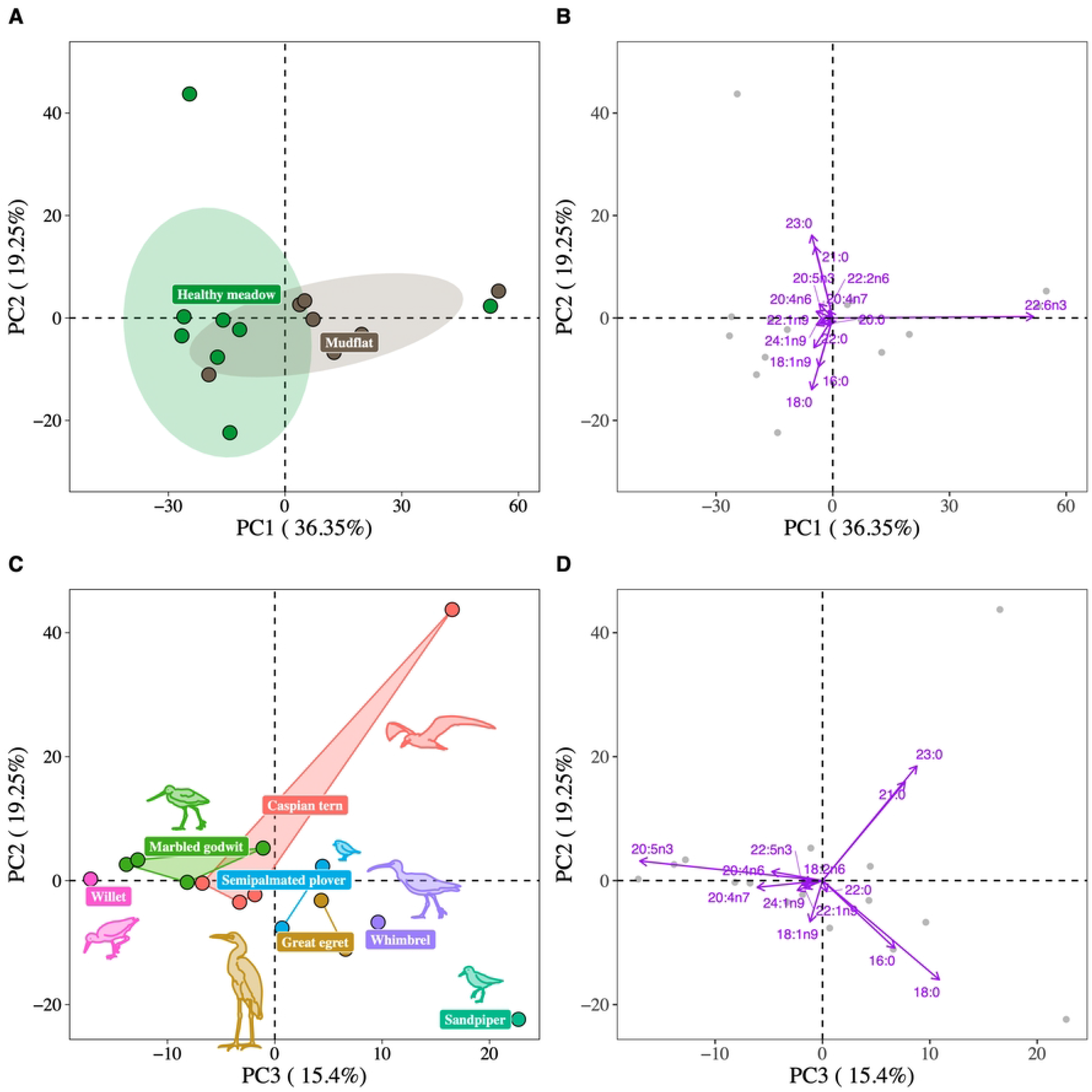
Free fatty acids (FFA) principal components analysis (PCA) results. Panel A) shows a Principal component (PC) 1 and PC2, with points colored by environment and ellipses drawn at a radius of 80%. Panel B) shows the eigenvectors for PC1 and PC2. Panel C) shows PC2 and PC3 with the points colored by bird species. Panel D) shows the eigenvectors for PC2 and PC3. For simplicity, only eigenvectors with cos^2^ > 0.6 are labelled in B) and D).

**Fig 4:**
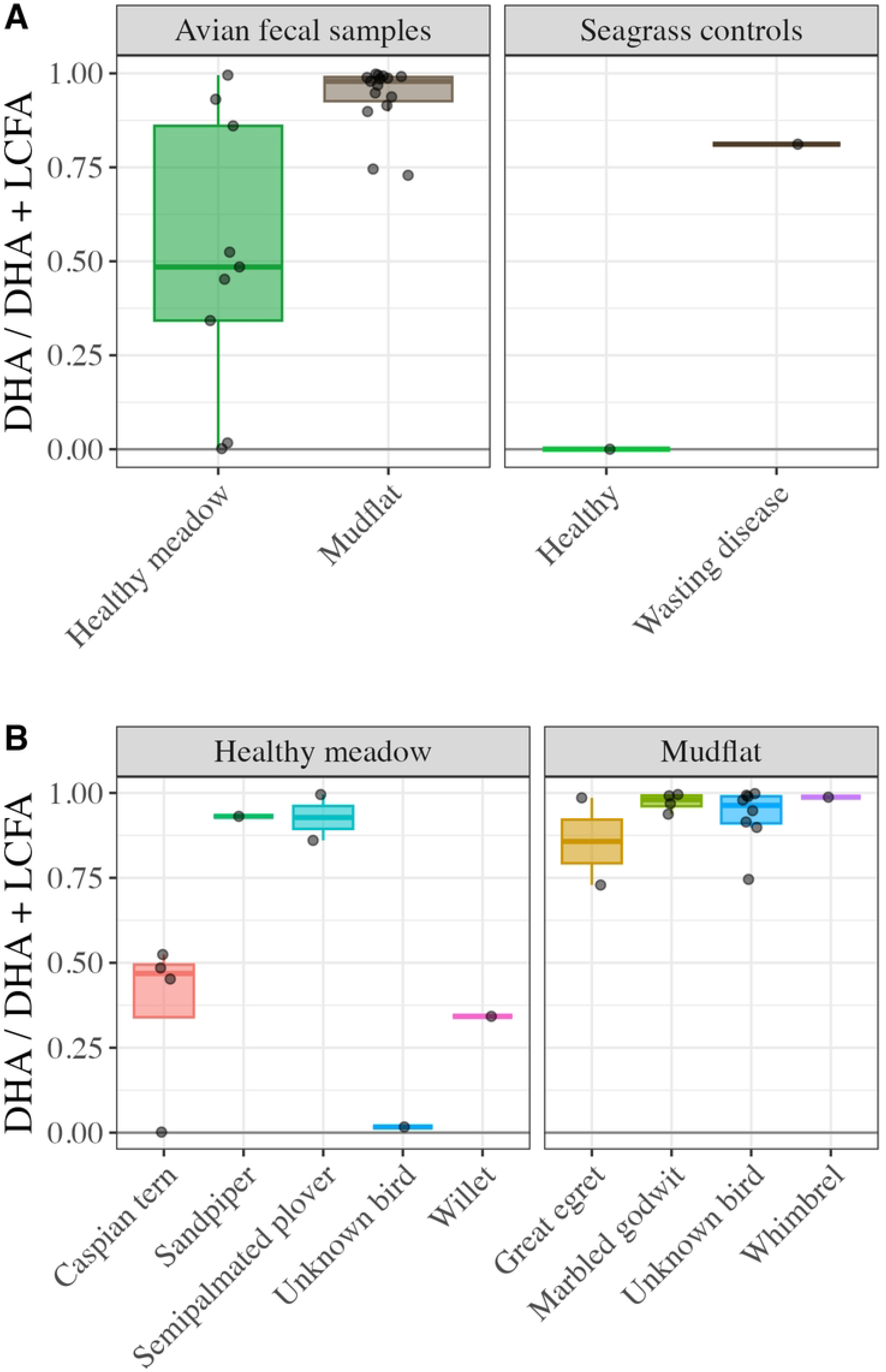
Barplots showing a ratio of docosahexaenoic acid (DHA) normalized to the total long-chain fatty acids (LCFAs (24:0 + 24:1n-9 + 25:0 + 26:0 + 28:0 + 29:0 + 30:0), including data from unknown species. Panel A) compares the two ecosystems and the two seagrass tissue controls. Panel B) shows the same information, isolating each bird species separated (including those that are unknown).

A total of eight ANOVA tests were performed between the two ecosystem types (see **Table 1**). Of these, only the DHA / (DHA + LCFA) differed significantly between ecosystems (p = 0.0155). An ANOVA of a linear regression for DHA / (DHA + LCFA) was done to validate the model, which returned an r^2^ value of 0.3734. The trends of each species were not tested with ANOVA since the sample sizes would violate the model assumptions. Animals are broadly efficient at metabolizing diverse FAs and at molecule-specific rates; it is therefore striking that we can correlate the fecal FA pattern to the ecosystem with such confidence.

## Discussion

While fecal samples from carnivores represent ecological black boxes with many unknown variables, the results of this pilot project suggest that they may be suitable for noninvasively monitoring ecosystems. Despite the limited dataset, fecal DHA normalized to LCFAs can distinguish between the healthy seagrass meadows and mudflats. It is possible that high LCFA values indicate terrestrial and freshwater input to the Millerton Point sample area, since it sits at the mouth of Lagunitas Creek. However, we can leverage the differential PUFA biosynthesis rates in terrestrial vs marine ecosystems to test for this bias (Colombo et al. 2017). The ratio of terrestrial/marine PUFAs, (18:2n-6 + 18:3n-3) / (20:4n-6 + 20:5n-3 + 22:6n-3), did not yield a significant ANOVA difference between the ecosystems (Table 1). Likewise, high DHA values could be indicative of Gaffney Point being closer to the ocean than Millerton Point, but these concerns are again assuaged by the PUFA ANOVA test, which shows no discernible bias in ecosystem inputs. It remains unclear what the primary source of DHA was, since it is produced by diverse organisms via *de novo* synthesis and modification of other FAs (Kharlamenko et al. 2001; Dalsgaard et al. 2003; Veloza et al. 2006; Chu et al. 2008; Coelho et al. 2011; Peltomaa et al. 2017; Yoshioka et al. 2019). Regardless of the specific source of DHA, the statistical significance holds, and it has been observed that healthy seagrass meadows can modulate their microbiomes to have much lower abundances of the microbial eukaryotes, especially diatoms and dinoflagellates (Jacobs-Palmer et al. 2020; Liu et al. 2022; Pan et al. 2022). The elevated presence of DHA in bird stool may suggest both the presence of the seagrass wasting disease and various microbial eukaryotes characteristic of a mudflat or collapsing meadow, and a low amount of DHA suggests a healthy seagrass meadow that is free of pathogens and/or healthy enough to remove diatoms and dinoflagellates from its microbiome.

The recovery of diverse, and presumably quickly metabolized, FFAs from bird stools lays the groundwork for broader research into avian biomonitoring techniques. Although it is a straightforward and common practice to identify host species from mammalian dung using DNA barcoding (such as 18S and COX1 genes), it is difficult to achieve the same results from bird fecal samples (Joo and Park 2012; Smith et al. 2023; Spence et al. 2025). The FFA approach here shows that lipid chemotaxonomy may offer a promising alternative for birds to traditional genomic barcoding methods when paired with fecal sample controls of known species. Moreover, FFAs from stool offer a snapshot of diet, which can be consumed and metabolized within ∼10-60 minutes for small birds depending on the species (Kuwae 2007). This represents a less invasive data source than typical FA samples taken from bird adipose tissue or blood plasma, as well as protocols that require euthanasia for samples of flight muscles, nervous system tissue, or stomach contents (Williams and Buck 2010). A larger implication of recovering essential FAs from bird stool is the previously unappreciated export of marine nutrients to terrestrial ecosystems via bird guano, in addition to classical elements such as nitrogen and phosphorus (Obrist et al. 2020; Appoo et al. 2024; Finne et al. 2024; Grant et al. 2024). Lastly, finding seagrass LCFAs in the food chain’s top carnivores highlights the nutritional value of intact seagrass meadows, due to the seagrass’s recalcitrant tissues it is likely that the LCFAs came from detritus rather than living biomass (Harrison 1989; Nichols et al. 1986; Kharlamenko et al. 2001).

While the results presented here are promising, there are many limitations to this study that need to be considered in future research. The field work was less efficient than traditional seagrass monitoring methods, especially with regard to the predictability of target bird species being present and the recovery of stool samples (∼100 hours of field work for 24 samples). While the birds may have very local chemical signatures, their prey–especially larger fish eaten by the terns and egrets–represent much longer averages of the environment and have potential to travel long distances to the field site. This concern compounds with the unknown number of organisms that could be in the trophic chain. Despite being a keystone species, not many organisms directly consume live seagrass, which restricts the mobilization of seagrass FAs amenable for proxy work to mostly detritus. This may make our approach prone to false negative results in locations with limited appropriate detritivores (Harrison 1989; Nichols et al. 1986; Kharlamenko et al. 2001).

An opposite way this could bias the results presented here is that the Tomales Bay seagrass prey community, specifically the Millerton Point area, is dominated by an invasive species of amphipod (*Ampithoe valida*) which directly consumes the seagrass blades and may be a driving force in the stronger seagrass signal than may be found elsewhere (Reynolds et al. 2012, Claire Murphy personal communication June 5^th^ 2026). Estimating a bird’s prey items from FAs is possible using Bayesian tools (such as the QFASA R package, Stock et al. 2018), which could help alleviate concerns about the ecosystem signal being biased by the species present. These advanced analyses need custom metabolic calibration coefficients and a strong local database - this is feasible to develop but would take extensive field work, prey taxa husbandry, and bird feeding trials to do properly (Iverson et al. 2004; Galloway and Budge 2020; Guerrero and Rogers 2020).

There are three main areas for future refinement for this monitoring technique: improved target species selection, larger datasets, and more cost-effective analyses. The efficiency of this new technique hinges largely on using species that can be predictably located, either by species (e.g. osprey nests and cormorant rookeries) or location (e.g. piers and buoys that birds tend to congregate around). Bird species that are too small or too terrestrial may not be as useful, as illustrated by the plover stool having a mudflat-type signal despite foraging in a seagrass-adjacent location. Picking better model species will require careful tradeoff consideration of their diet, foraging habits, and gut retention times. Validating this work will require expanded datasets, both temporally (e.g. sampling throughout the disease progression cycle over a summer) or spatially (e.g. along a long shoreline that encompasses multiple ecosystem gradients). Pairing this approach with collaborators who can diagnose disease severity indices could generate a scale of wasting disease rather than mere presence/absence. The addition of isotope work would greatly improve our understanding of the ecological interactions, especially δ^15^N and δ^13^C, which are used to estimate food chain length, and compound-specific isotope analysis on individual DHA molecules could disentangle their sources (Perkins et al. 2014; Dirir et al. 2025). While there is no equal substitute for mass spectrometry techniques for quantitative data, thin layer chromatography (TLC) or other gel electrophoresis techniques may offer a cheaper and easier qualitative method to identify target compounds now that their presence has been verified. The next steps for validating this lipid biomonitoring work will sample stools across the seagrass’s growing season from several locations and targeting species that congregate at specific locations, such as cormorants and osprey.

The pilot study described here is the basis to developing a noninvasive tool that could be readily implemented into existing infrastructure (e.g. birding groups and established environmental study programs), which can both complement and augment current seagrass monitoring practices. Developing efficient tools for monitoring seagrass meadow health, and therefore its carbon sequestration capacity, is imperative for international climate change initiatives and meadow restoration efforts (Unsworth et al. 2022; Cooley et al. 2022; Briand et al. 2026).

Ecosystem connectivity provides novel methods for monitoring keystone species beyond just seagrass - lipid proxies for autotrophs collected from carnivores allow researchers to pull on proverbial strings in a food web, affording us both a ‘top-down’ and ‘bottom-up’ views of the system as a whole when combined with traditional practices (Marneweck et al. 2022; Natsukawa and Sergio 2022; Briand et al. 2026). Many carbon dynamics are mediated by animals, especially carnivores like birds that can travel between different ecosystems; lipidomics is a powerful tool to efficiently reconstruct these endmember relationships (Schmitz et al. 2010; Schmitz et al. 2014; Buelow and Sheaves 2015). Once better validated, this new technique could be expanded to any inaccessible ecosystem with a keystone species that has unique lipids and a mobile animal that goes in and back out to somewhere more accessible. For example, kelp forests are incredibly important and imperiled, but scuba-diving monitoring and restoration programs in California are often hindered by the white sharks that hunt there. Other potential ecosystems to monitor could include coral reefs, mangroves, and deep-sea ecosystems. This animal-based lipid biomonitoring technique is a versatile and holistic approach to conservation biology: by tracing specific molecules up through ecosystems, an animal’s natural behavior can bring samples directly to researchers.

## Author Contribution Statement

Christopher Mulligan: Conceptualization, Funding acquisition, Methodology, Resources, Investigation, Data curation, Visualization, Formal Analysis, Writing - original draft, Writing - review & editing. David A. Gold: Resources, Supervision, Writing - review & editing.

## Data accessibility statement

The data that support the findings of this study are openly available on Github at https://github.com/DavidGoldLab/2026_Seagrass_Monitoring.

## Acknowledgements

This work was made possible by many wonderful people: Alyssa Griffin, Nils Warnock, and Jackie Sones gave advice and helped with fieldwork logistics; Serina Moheed gave field site advice and provided the seagrass tissue controls; Philip Barruel helped with HPAI safety; Sarah King, Anjelica Guerrier, and Howard Higley were fieldwork assistants; and Jay Stachowicz and Aaron Galloway gave valuable feedback on the results.

## Supporting information

**S1 Fig. PCA of lipids across the two field sites.** This includes all lipids, not just free fatty acids. It also includes compounds such as sterols and triacylglycerides.

**S1 Table. UHP-LC data set.** This provides the full 4,067 compound UHP-LC dataset for all samples, including molecular mass and annotation information.

**S2 Table. Fatty acid data set.** The free fatty acid values for each sample used in the analyses.

